# Increasing the Spatial Extent of Attention Strengthens Surround Suppression

**DOI:** 10.1101/2021.11.26.470072

**Authors:** Merve Kiniklioglu, Huseyin Boyaci

**Affiliations:** Interdisciplinary Neuroscience Program, Bilkent University, Ankara, 06800, Turkey; Aysel Sabuncu Brain Research Center & National Magnetic Resonance Research Center (UMRAM), Bilkent University, Ankara, 06800, Turkey; Department of Psychology, Bilkent University, Ankara, 06800, Turkey; Department of Psychology, Justus Liebig University Giessen, Giessen, Germany

**Keywords:** attention, center-surround interaction, motion perception, response normalization, surround suppression

## Abstract

Here we investigate how the extent of spatial attention affects center-surround interaction in visual motion processing. To do so, we measured motion direction discrimination thresholds in humans using drifting gratings and two attention conditions. Under the narrow attention condition, attention was limited to the central part of the visual stimulus, whereas under the wide attention condition, it was directed to both the center and surround of the stimulus. We found stronger surround suppression under the wide attention condition. The magnitude of the attention effect increased with the size of the surround when the stimulus had low contrast, but did not change when it had high contrast. Results also showed that attention had a weaker effect when the center and surround gratings drifted in opposite directions. Next, to establish a link between the behavioral results and the neuronal response characteristics, we performed computer simulations using the divisive normalization model. Our simulations showed that the model can successfully predict the observed behavioral results using parameters derived from the medial temporal (MT) area of the cortex. These findings reveal the critical role of spatial attention on surround suppression and establish a link between neuronal activity and behavior. Further, these results also suggest that the reduced surround suppression found in certain clinical disorders (e.g., schizophrenia and autism spectrum disorder) may be caused by abnormal attention mechanisms.

## 1 Introduction

Sensitivity to a visual stimulus often depends on what is presented in its surround. For example, contrast detection thresholds increase if a grating is surrounded by iso-oriented high-contrast stimuli (Petrov, Carandini, & McKee, 2005; Xing & Heeger, 2001). Motion direction discrimination thresholds are also affected by these ‘center-surround’ interactions. As the size of a drifting grating increases, direction discrimination thresholds increase for high contrast, and decrease for low contrast stimuli (Tadin, Lappin, Gilroy, & Blake, 2003). In line with the behavioral results, a neuron’s responses to a stimulus presented at the center of its receptive field is suppressed by iso-oriented stimuli presented in its surround (Cavanaugh, Bair, & Movshon, 2002a; Shushruth et al., 2012). This kind of surround suppression in neurons is observed for both static and dynamic stimuli at many levels of the visual system (Angelucci et al., 2017), including the primary visual cortex (V1), and middle temporal complex (hMT) (Er, Pamir, & Boyaci, 2020; Schallmo et al., 2018; Turkozer, Pamir, & Boyaci, 2016). Thus surround suppression at the neuronal level is believed to shape visual sensitivity at the behavioral level for static (Shushruth et al., 2013) as well as dynamic stimuli (Tadin, 2015).

Surround suppression is known to be critically affected by spatial attention (Maunsell, 2015). For example, in non-human animals directing attention to the center of a static stimulus results in weaker neuronal suppression compared to directing attention to the surround (Reynolds & Chelazzi, 2004; Sundberg, Mitchell, & Reynolds, 2009; also see Ito & Gilbert, 1999; Roberts, Delicato, Herrero, Gieselmann, & Thiele, 2007). The effect of spatial attention using static stimuli, both behaviorally and neuronally, is documented in humans, as well (Flevaris & Murray, 2015a, 2015b; Freeman, Sagi, & Driver, 2001; Herrmann, Montaser-Kouhsari, Carrasco, & Heeger, 2010). Dynamic stimuli, on the other hand, is used in only a few non-human animal studies to assess the effect of attention on surround suppression (Anton-Erxleben, Stephan, & Treue, 2009; Womelsdorf, Anton-Erxleben, Pieper, & Treue, 2006). No previous study on humans, to the best of our knowledge, has systematically examined the effect of attention on surround suppression using motion stimuli.

Indirectly, however, there is evidence that attention may modulate surround suppression with motion stimuli in humans. Using drifting gratings, autism spectrum disorder (ASD) patients have been found to exhibit weaker surround suppression compared to a neurotypical group (Schallmo et al., 2020). The authors have argued that the weak suppression can be explained by a narrow spatial attention field, and supported their argument with computer simulations using the divisive normalization model of attention (Reynolds & Heeger, 2009).

Divisive normalization model can successfully predict the effects of attention on neuronal and behavioral responses. The model suggests that attention scales the excitatory and inhibitory drives received by a neuron, which together determine the overall firing rate of that neuron. Accordingly, the spatial spread of attention field affects the contrast-response functions of neurons, and consequently, behavioral sensitivity. Specifically, the model predicts that a narrower attention field causes weaker suppression because it primarily increases the effect of the excitatory drive. Conversely, a wider attention field leads to stronger suppression because it increases both the effects of excitatory and inhibitory drives.

Here, we investigated the effect of spatial extent of attention on surround suppression in humans using motion stimuli. We first conducted a behavioral experiment with two attention conditions: narrow attention condition in which attention is restricted to the center of the stimulus, and wide attention condition in which attention is directed to both the center and surround. If attention had an effect, we would expect stronger suppression under the wide attention condition compared to narrow attention condition. To anticipate, we found that surround suppression is stronger under the wide attention condition. Next, by using the divisive normalization model and incorporating the spatial extent of attention, we sought to establish a link between the behavioral results and neuronal response characteristics.

## 2 Methods

### 2.1 Participants

Ten female volunteers, (mean age= 26.7 years), participated in the experiment. All participants reported normal or corrected-to-normal vision and had no history of neurological or visual disorders. Participants gave their written informed consent before the experiment. The experimental protocols were approved by the Human Ethics Committee of Bilkent University.

### 2.2 Apparatus

Stimuli were generated using the Psychophysics toolbox (Brainard, 1997) with MATLAB 2018a (MathWorks, Natick, MA) and presented on a CRT monitor (HP P1230, 22 inches, 1280 × 1024 resolution, refresh rate 120 Hz). Participants were seated in a dark room with their heads stabilized using a chin rest at a distance of 75 cm from the monitor. A gray-scale look-up table was prepared through direct measurements of the luminance values (SpectroCAL, Cambridge Research Systems Ltd., UK), and used to ensure the presentation of correct luminance values. Responses were collected via a standard computer keyboard.

### 2.3 Stimuli and Design

Stimuli were vertically oriented drifting sinusoidal gratings (spatial frequency: 1 cycle per degree) weighted by two-dimensional raised cosine envelopes (Foss-Feig, Tadin, Schauder, & Cascio, 2013), the radius of which defined the stimulus size. Specifically, the stimulus consisted of a center grating surrounded by an annular grating (except the center-only configuration under the narrow attention condition, see below), which drifted within the stationary raised cosine envelopes (starting phase randomized) at a rate of 4°/s either leftward or rightward (Figure 1). The stimuli were generated using built-in functions of Psychtoolbox, and presented foveally on a mid-gray background (16.09 cd/m^2^).

**Figure 1:**
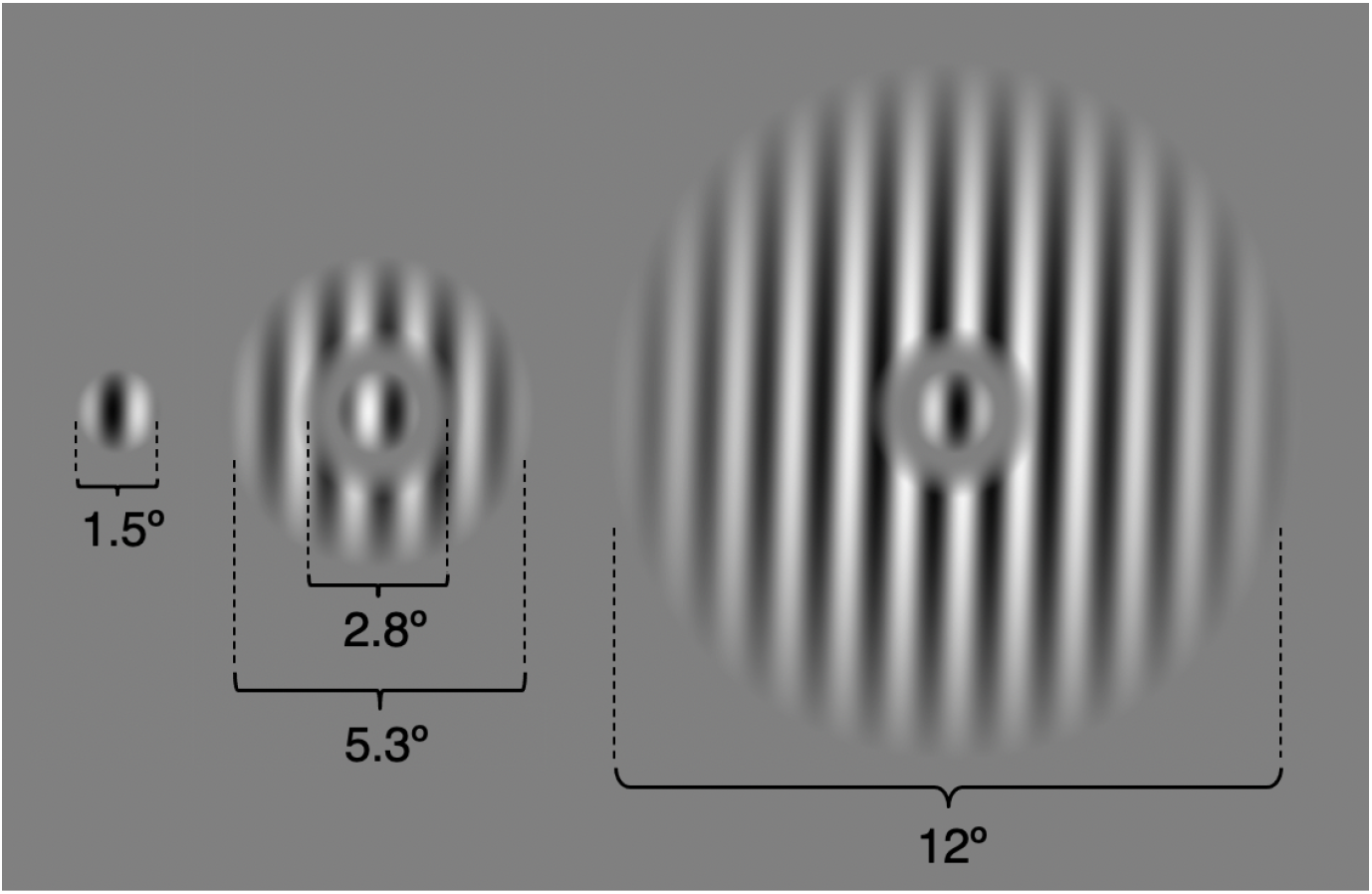
Center-only, small-surround, and large-surround configurations for high (98%) contrast stimuli.

In the narrow attention condition, a center sinusoidal grating is either presented alone or surrounded by a surround grating. Trials in which the center stimulus is presented alone (center-only) were used as a baseline to calculate the suppressive effect of the surround on the center. In the wide attention condition, the stimulus consisted of a center sinusoidal grating surrounded by a surround grating. The diameter of the center grating was 1.5°. The surround gratings were presented in two different sizes. In the small-surround configuration, the inner diameter of the surround was 2.8° and its outer diameter was 5.3°. In the large-surround configuration, the inner diameter of the surround was 2.8° and its outer diameter was 12°. The area between (1.5° to 2.8°) remained unstimulated to separate the center grating from the surround grating (Figure 1).

The motion directions of the center and surround gratings were randomly determined for each trial and were equally probable in either direction. In half of the trials the center and surround gratings moved in the same direction, in the other half in opposite directions.

There were two contrast conditions in the experiment. The Michelson contrast of both center and surround gratings were 3% in the low contrast condition, and 98% in the high contrast condition. Two conditions of attention (narrow attention and wide attention), and two contrast levels (3% and 98%) were tested in separate sessions (4 experimental sessions in total) whereas two size levels (small-surround and large-surround) and two motion direction (same-direction and opposite-direction) were tested in the same session in randomized order. The order of the sessions was counterbalanced for each participant and scheduled on different days.

For each trial, participants viewed foveally presented drifting gratings and performed a task about the perceived direction of motion via a keypress. In the narrow attention condition, participants were asked to attend the center grating and report its motion direction. In the wide attention condition, participants were instructed to attend both to the center and surround gratings. They first reported the drift direction of the center grating, then reported whether the center and surround gratings drifted in the same direction. This was done solely to encourage the participants to widen their attention fields. Hence, any difference in motion direction discrimination thresholds between narrow and wide attention conditions must reflect the effect of the spatial extent of attention as the stimulus presented in those conditions were identical (Figure 2).

**Figure 2:**
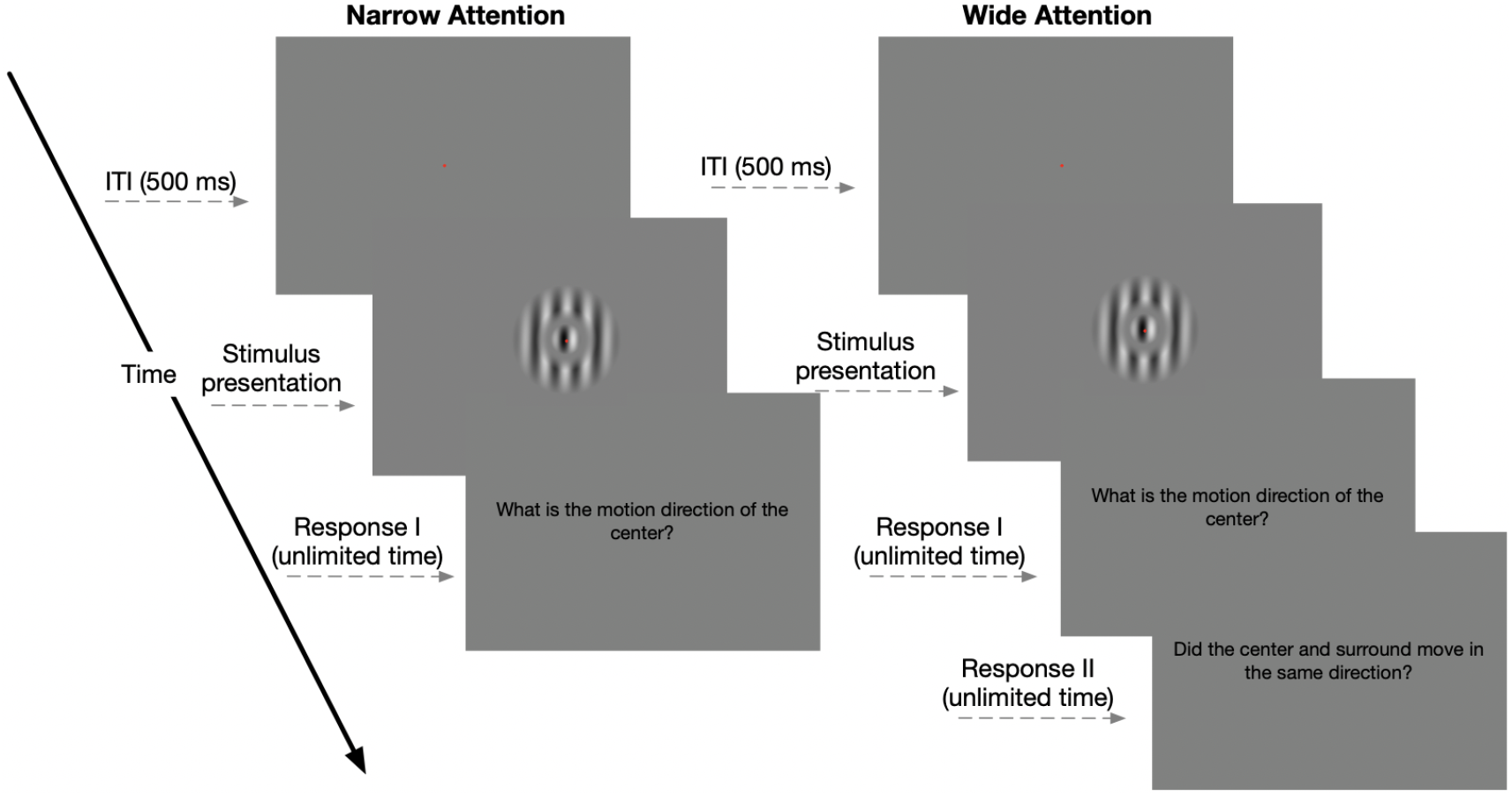
Experimental design for the narrow and wide attention conditions. Participants were asked to keep their eyes on a fixation point throughout the trial. Each trial began with a fixation point followed by the presentation of the stimulus, whose duration was adjusted with two interleaved 1-up 3-down staircase. In the narrow attention condition, participants reported the motion direction of the center grating. In the wide attention condition, participants first reported the drift direction of the central grating, then reported whether the central and annular gratings drifted in the same direction.

In each trial of all conditions, the duration of the presentation, defined as 2 standard deviation (SD) of a temporal Gaussian envelope (Borghuis, Tadin, Lankheet, Lappin, & van de Grind, 2019; Tadin et al., 2003), was adjusted following an adaptive procedure (two interleaved 1-up 3-down staircases), based on the participants’ judgments of the center grating drift direction in previous trials. Importantly, participants’ judgments about the motion direction of the surround are not used for adaptively changing the stimulus duration; this second task is only used to encourage them to extend their attention field. There were two independently progressing staircases for each condition. One staircase started from a very short duration (25 ms) which made the task relatively harder, the other started from a long duration (158 ms), which made the task relatively easier. There were 120 trials in each staircase. Each subject completed 1200 trials for narrow attention and 960 trials for wide attention conditions for each contrast. Each session took approximately 45 minutes. Brief break periods were given during the experiment.

### 2.4 Data Analysis

Duration thresholds (79% success rate) were calculated by fitting the responses with a Weibull function using the Palamedes toolbox (Kingdom & Prins, 2010) in MATLAB 2019a (MathWorks, Natick, MA) for each participant and condition. Next, using the threshold values we calculated Suppression Indexes (SI) to quantify the strength of the surround suppression

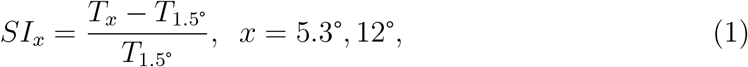

where *T*_._ is the discrimination threshold for a given size. Higher positive values of SI indicate stronger surround suppression, whereas negative SI values mean surround facilitation. An SI of 0 means no suppression or facilitation.

We compared the SI values to “0” by applying a one-sample two-tailed Student’s t-test with correction for multiple tests using SPSS Version 25 (SPSS Inc., Chicago, IL) to observe the expected effect of suppression. Next, we performed three-way repeated measures ANOVA on same-direction trials with three factors: attention (narrow- and wide-attention), size (small- and large-surround), and contrast (high- and low-contrast) to explore how the size-contrast interaction is affected by the spatial extent of the attention.

Then, we conducted pre-planned t-tests to test our research questions. Firstly, *is the surround suppression stronger in the wide attention condition compared to the narrow attention condition?* To find an answer to this question we tested whether the mean *SI* under the wide attention condition is significantly larger than that under the narrow attention condition. Secondly, *does surround suppression increase with the size of the surround stimulus more strongly in the wide attention condition compared to narrow attention condition?* To find an answer to this question, we tested whether the slope of the rise in SI with the size is steeper under the wide attention condition compared to the narrow attention condition by using two-tailed paired-samples t-test. Finally, *if there are these effects, are they purely because attention is allocated to a wider region, or is it because attention modulates surround suppression?* To answer this question and make sure that the observed effect is not a task-demand artifact, we performed two-tailed paired-samples Student’s t-test to test whether the effects of attention are stronger under the same-direction condition than the opposite-direction condition.

### 2.5 Normalization Model

We used the normalization model of attention (NMA) (Reynolds & Heeger, 2009) to establish a link between the behavioral results and possible neuronal response characteristics. The model simulates the response of a population of neurons that are tuned to spatial position (spatial attention) and motion direction (feature-based attention). The model has three components:

1. Stimulus Drive, which represents the excitatory response of a neuron in the absence of suppression or attention;
2. Suppressive Drive, which represents summed activity of a pool of neurons;
3. Attention Field, which modulates attentional gain for each neuron in the population according to spatial position and direction tuning (see Figure 3).

**Figure 3:**
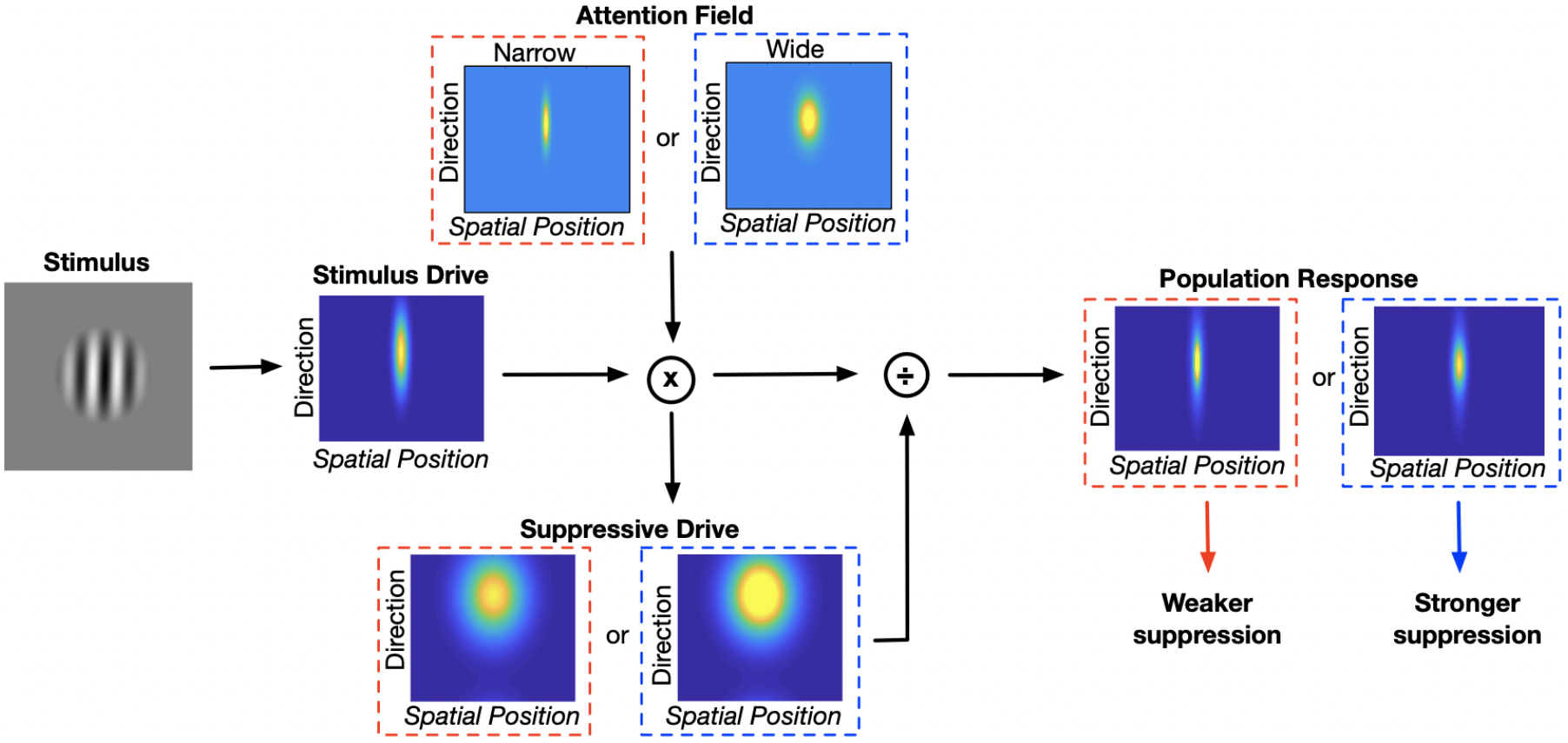
A schematic representation of the normalization model of attention. Stimulus Drive is multiplied by the attention field, then normalized by the Suppressive Drive to determine population response. We simulated model responses using a narrow attention field for the narrow attention condition (red dashed squares), and a wider attention field for the wide attention condition (blue dashed squares) (Eqs. 2–4).

To determine the population response the stimulus drive is multiplied by the attention field and then divided by the suppressive drive. The model can be summarized by the following equation:

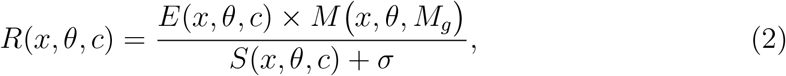

where *R* is predicted population response as a function of spatial position (*x*), direction (*θ*), and contrast (*c*). *E* and *S* are the excitatory and suppressive drives respectively, and *σ* is the semi-saturation constant, which prevents *R* from being undefined when *S* equals zero. Excitatory drive, *E*, is computed by the following equation:

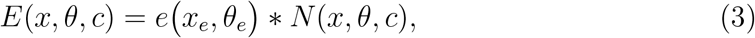

where *e* is a two-dimensional (2-D) Gaussian function with the spatial position (*x_e_*) and the direction (*θ_e_*) that determines spatial and direction tuning, respectively for the excitatory drive (*E*), ∗ denotes convolution, and *N* is the neural representation of the stimulus, which is also a 2-dimensional Gaussian function with the stimulus position (*x*), and the direction (*θ*), and scaled by the contrast (*c*). Since our stimulus consisted of a center grating surrounded by a surround grating, *N* is obtained by the summation of the two 2-D Gaussian functions that represent the stimulus image for the center grating and the surround grating.

Suppressive drive, *S*, is computed by the following equation:

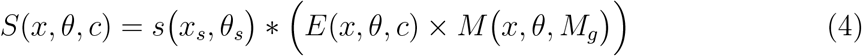

where *s* is another 2-D Gaussian function as *e*, but shows broader tuning in spatial position (*x_e_*) and direction (*θ_s_*). *M* represents the attention field, which reflects how spatial attention modulates the stimulus processing by scaling *E*. *M* is a 2-D Gaussian function, whose spatial extent is set by the attentional gain factor *M_g_*. The operator “×” represents element-wise multiplication (Hadamard product).

We computed predicted motion discrimination thresholds from model responses as

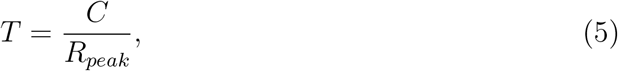

where *R_peak_* is the maximum of the predicted model response *R* (winner-take-all approach, see Er et al., 2020; Schallmo et al., 2020) and *C* is a constant linking the thresholds and neuronal responses (Schallmo et al., 2020).

Because our stimulus was consisted of a center grating surrounded by an annular grating, we modeled neural response to this stimulus by adding neural representation of center and surround stimulus. We summed two different 2-D Gaussian functions (*N*, see Equation 3) corresponding to center and annular gratings. To model the neural responses to the narrow and wide attention conditions, we used a smaller (*M_g_*, 3 arbitrary units) and larger (*M_g_*, 9 arbitrary units) spatial attention width, respectively. Moreover, the opposite direction trials are modeled by using surround stimulus direction that is 180° away from the direction of the center. To be more specific, the stimulus direction parameter for the 2-D Gaussian function was −90° for the center and 90° for the surround (see e.g., Reynolds & Heeger, 2009).

Table 1 reports the parameters used in simulations that were choosen based on the MT neurons’ receptive field properties reported in the literature (Schallmo et al., 2018, 2020). Simulations were done on Matlab 2018, separately for same-direction and opposite-direction trials. The model was implemented using custom MATLAB functions (Er et al., 2020; Schallmo et al., 2020).

**Table 1:**
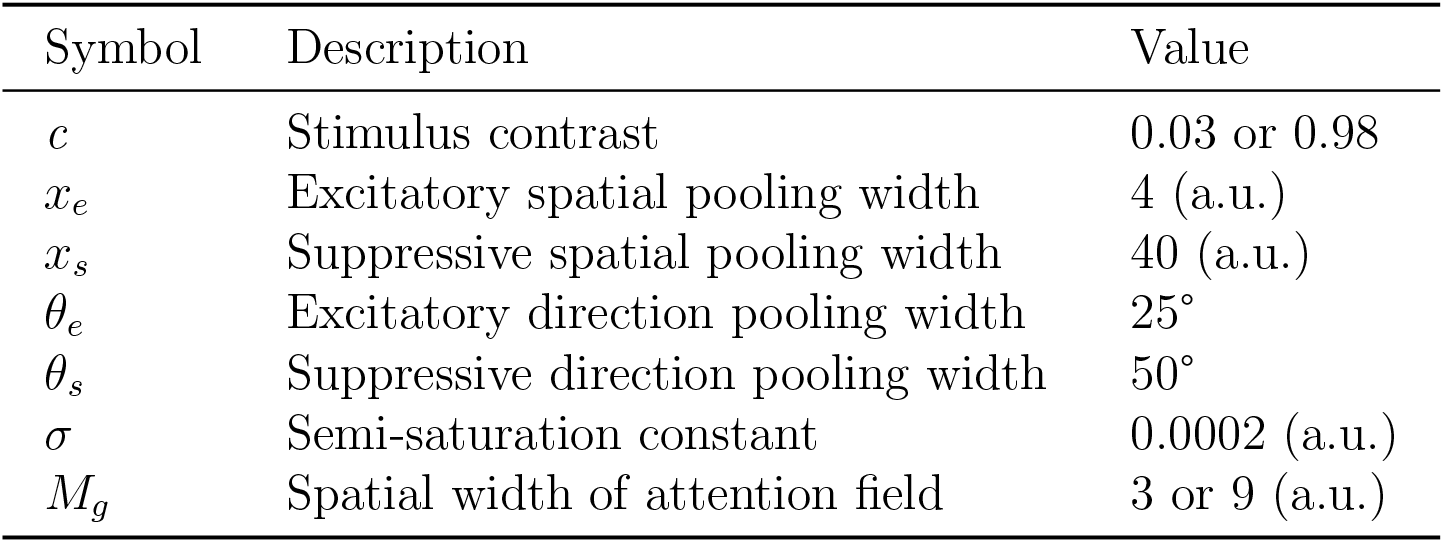
Normalization model parameters.

| Symbol | Description | Value |
| --- | --- | --- |
| $c$ | Stimulus contrast | 0.03 or 0.98 |
| $x_e$ | Excitatory spatial pooling width | 4 (a.u.) |
| $x_s$ | Suppressive spatial pooling width | 40 (a.u.) |
| $\theta_e$ | Excitatory direction pooling width | $25^\circ$ |
| $\theta_s$ | Suppressive direction pooling width | $50^\circ$ |
| $\sigma$ | Semi-saturation constant | 0.0002 (a.u.) |
| $M_g$ | Spatial width of attention field | 3 or 9 (a.u.) |

## 3 Results

### 3.1 Behavioral Results

Figure 4 shows motion direction discrimination thresholds averaged across all ten participants, and Figure 5 shows suppression indexes (*SI*, Methods) derived from the threshold values. Results of same direction trials (Figs. 4–5, A&B) exhibit a pattern that is consistent with literature (Tadin et al., 2003), especially for the high contrast condition. Firstly, under both attention conditions *SI* values increase with size. Secondly, *SI* values are significantly larger than zero for all conditions except the low-contrast small-surround trials under both the narrow and wide attention conditions, which shows that for a low-contrast stimulus increase in size up to a value may lead to surround facilitation (Er et al., 2020; Schallmo et al., 2018; Tadin et al., 2003). Table 2 shows one-sample *t*-test results.

**Figure 4:**
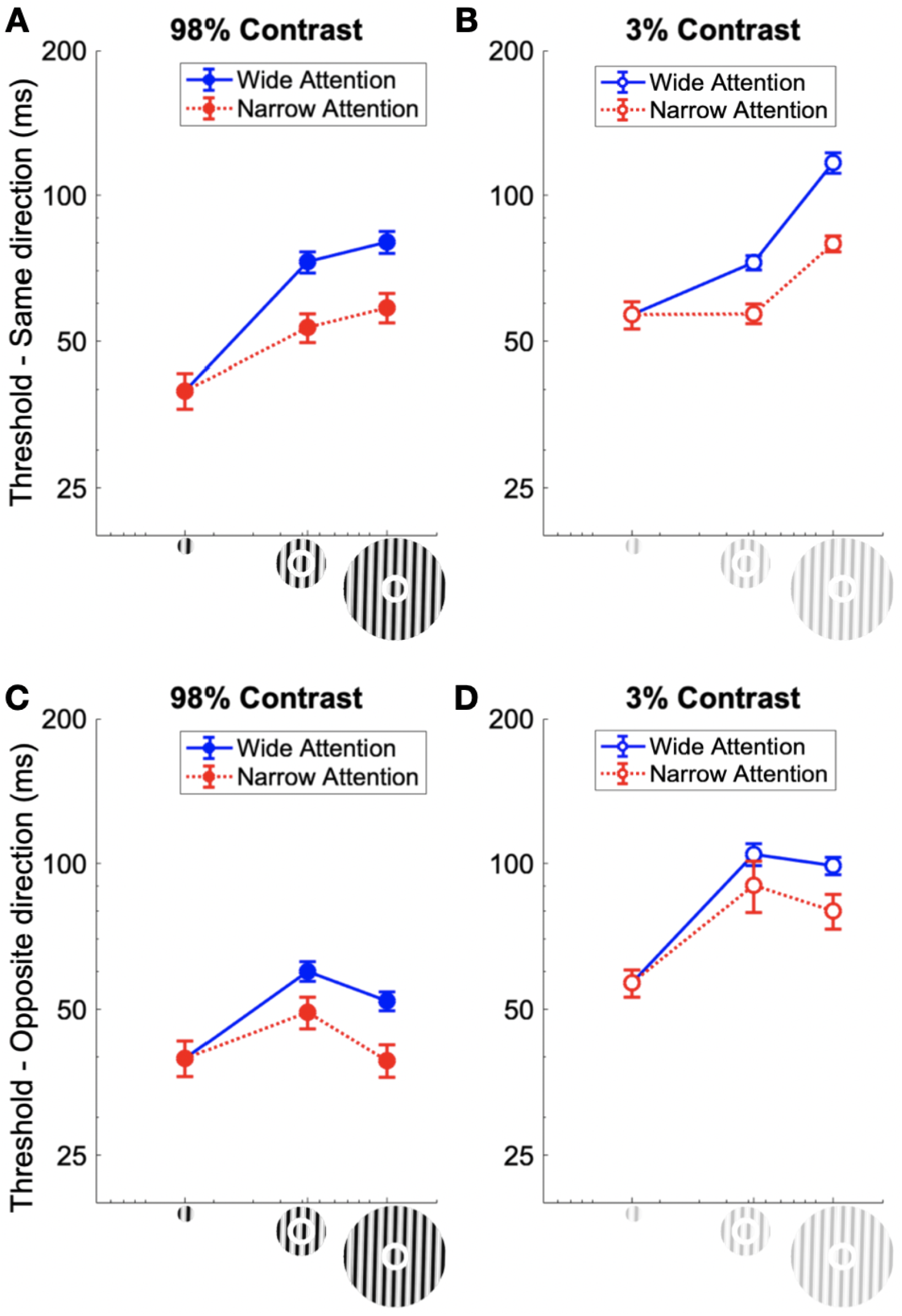
Behavioral results. Motion direction discrimination thresholds averaged across participants. A and B for same direction trials, C and D for opposite direction trials. Error bars represent SEM.

**Figure 5:**
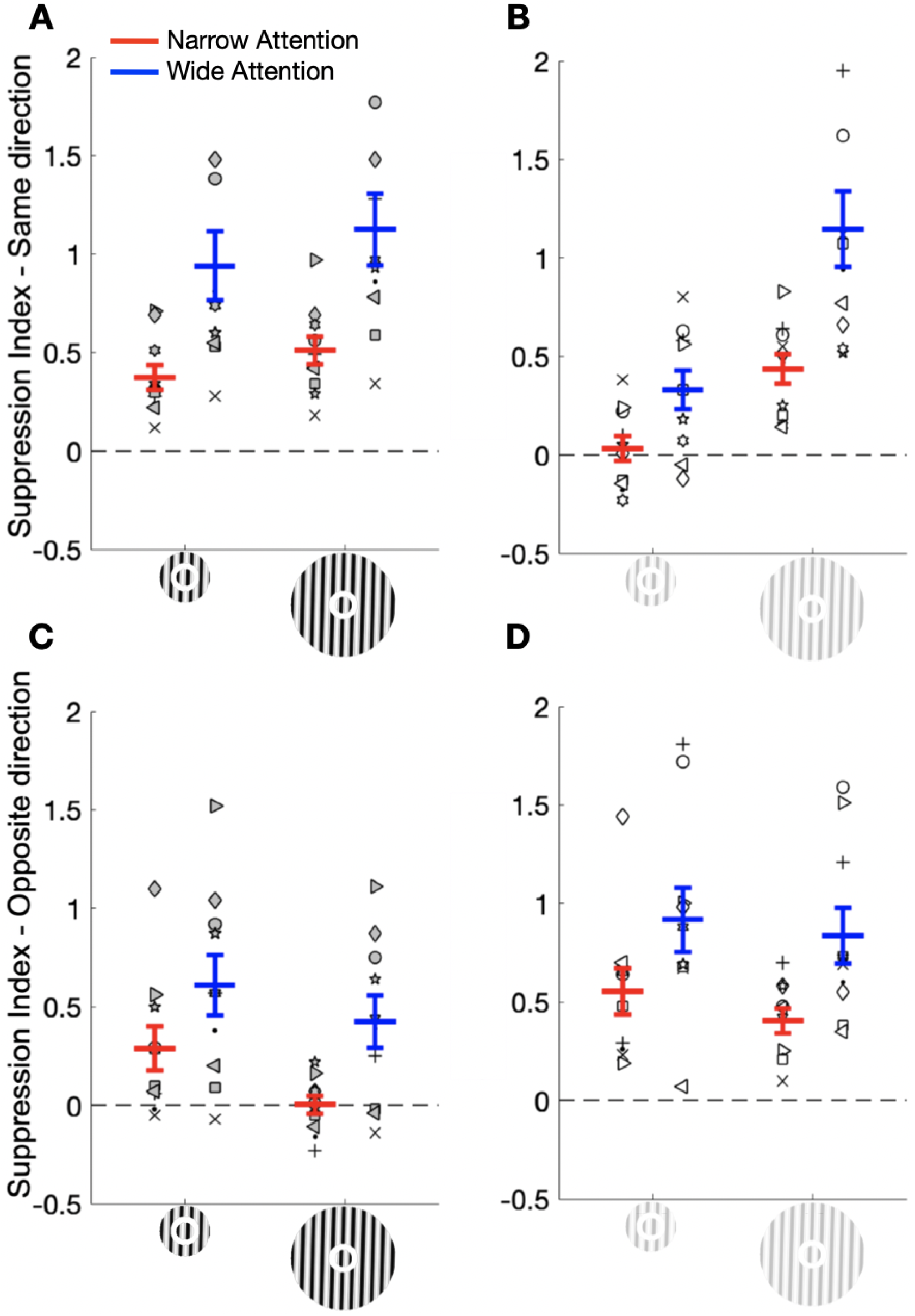
Supression Indices (SI). A and B, same direction trials, C and D, opposite direction trials. Higher values of *SI* indicate stronger surround suppression, negative *SI* values indicate surround facilitation. Symbols represent individual participants, and the red and blue lines show averages for narrow and wide attention conditions, respectively. Error bars represent SEM.

**Table 2:** Results of one-sample *t*-test for same-direction trials (*H*_0_ : *SI* = 0). Numbers in degrees are the surround diameters.

| | Mean Difference | $df$ | $t$ | $p$ |
| --- | --- | --- | --- | --- |
| Same Direction |  |  |  |  |
| Narrow Low 5.3° | 0.03 | 9 | 0.50 | 0.63 |
| Narrow Low 12° | 0.44 | 9 | 5.76 | < 0.001* |
| Narrow High 5.3° | 0.37 | 9 | 5.94 | < 0.001* |
| Narrow High 12° | 0.51 | 9 | 7.17 | < 0.001* |
| Wide Low 5.3° | 0.33 | 9 | 3.41 | 0.008 |
| Wide Low 12° | 1.15 | 9 | 5.94 | < 0.001* |
| Wide High 5.3° | 0.94 | 9 | 5.39 | < 0.001* |
| Wide High 12° | 1.13 | 9 | 6.16 | < 0.001* |

$df$ = Degrees of Freedom, \* indicates significance at $\alpha_{corr} = 0.00625$ .

More importantly, in all conditions we visually observe stronger surround suppression under the wide compared to narrow attention condition (Figs. 4, 5). Robustness of this observation is supported by three-way repeated measures ANOVA results (Table 3), which showed significant main effects of attention and size, as well as size-contrast, attention-size and attention-size-contrast interactions.

**Table 3:** Results of three-way ANOVA for same direction trials.

| Source | $SS$ | $F(1, 9)$ | $p$ | $\eta^2$ |
| --- | --- | --- | --- | --- |
| Same Direction |  |  |  |  |
| attention | 6 | 35.97 | <0.001 | 0.80 |
| Contrast | 1.25 | 4.75 | 0.057 | 0.35 |
| Size | 2.98 | 50.76 | <0.001 | 0.85 |
| attention × contrast | 0.04 | 0.49 | 0.50 | 0.05 |
| attention × size | 0.26 | 13.22 | 0.005 | 0.60 |
| Contrast × size | 1 | 31.32 | <0.001 | 0.78 |
| attention × contrast × size | 0.17 | 8.13 | 0.019 | 0.48 |

$SS$ = Sum of Squares

Next we focus on our pre-planned comparisons. Our first comparison suggests that increasing the spatial extent of attention evokes stronger suppression: *SI* was significantly larger in the wide compared to the narrow attention condition (main effect of attention; *F*_(1,9)_ = 35.97, *p* < 0.001), and for all individual conditions, *SI* values were significantly higher in the wide attention condition compared to narrow attention condition (*p*s < 0.002, *α_corr_*=0.0125). Further, *SI* increased with the size of surround more strongly in the wide compared to the narrow attention condition (*t*_(9)_ = 3.64, *p* = 0.005). These provide answers to our first two research questions. Namely they show that surround suppression is stronger in the wide attention condition compared to the narrow attention condition, and surround suppression increases with the size of the surround stimulus more strongly in the wide attention condition compared to narrow attention condition.

It could be, however, argued that the results we found are just due to the demand of the second task (*i.e.* a task-demand artifact), and they do not reflect the effects of surround suppression mechanisms per se. To control for this possible confound, we compared the results of same-direction and opposite-direction trials. We did this because it is well known that at the single-unit level surround suppression is reduced or eliminated when surround and center move in opposite directions (Allman, Miezin, & McGuinness, 1985; Born & Tootell, 1992; Cavanaugh, Bair, & Movshon, 2002b; Kastner, Nothdurft, & Pigarev, 1995; Lamme, 1995). Thus we reasoned that if our results reflect a true effect of attention on surround suppression, then the difference in *SI* between wide and narrow attention conditions should be smaller in the opposite direction trials compared to those in the same direction trials. The patterns shown in Figure 5C and D for opposite-direction trials are clearly different than those in the same-direction trials shown in Figure 5A and B. To confirm, we applied two-tailed paired-samples Student’s *t*-test to see whether the difference in *SI* values between the narrow and wide attention condition was significantly larger in the same-direction trials compared to opposite-direction trials. Results showed that the effect of attention (as quantified by the *SI* difference between narrow and wide attention conditions) was significantly stronger in the same-direction compared to the opposite-direction trials (*t*_(9)_ = 3.11, *p* = 0.013). These results suggest that attention had a weaker effect on the thresholds when the center and surround gratings drifted in opposite directions. This answers our third research question, namely that the effect found in same-direction trials is not an artifact of task-demand, it reflects a genuine interaction between attention field and surround suppression.

Finally, we performed a post-hoc test to further explore whether attention interacts with size differently depending on the contrast of the stimulus. For low contrast, the difference in SI values between narrow and wide attention conditions in small-surround configuration is significantly smaller than that in large-surround configuration (*t*_(9)_ = 3.53, *p* = 0.006; *a_corr_* = 0.025). However, for high contrast, the effect of attention was identical in different size conditions (*t*_(9)_ = 0.96, *p* = 0.36). This shows that the magnitude of attention effect increases with the surround size in low contrast, but does not change in high contrast condition.

### 3.2 Model Results

We next sought to establish a link between our behavioral results and possible neuronal mechanisms. For this purpose we implemented a model based on the divisive normalization model incorporating spatial attention (Carandini & Heeger, 2012; Heeger, 1992). Critically, we used parameters that characterize MT neurons (Methods), because previous literature suggests that the neuronal activity in (h)MT area can successfully explain the size-contrast interaction in motion discrimination, both at the behavioral (Schallmo et al., 2018, 2020) and neuronal levels (Er et al., 2020; Herrmann et al., 2010; Lee & Maunsell, 2010b; Schallmo et al., 2018). To simulate the trials under the wide and narrow attention conditions we simply used a wider and a narrower attention field, respectively (Figure 3, Methods).

Although feature-based attention was not the focus of the current study, same versus opposite direction of motion incidentally introduce feature-based attention as a factor in our design. In the normalization model of attention, attention field is not only selective for a specific position (spatial attention) but also for direction of motion (feature-based attention). Hence, to address this possible effect, we included a feature-based attention component in the model (Figure 3, Methods).

Figure 6 shows the simulation results. These results are qualitatively similar to the behavioral findings shown in Figure 4. Specifically, simulated thresholds are lower with small attention field compared to those with large attention field, which is consistent with the weaker surround suppression under the narrow compared to wide attention condition that we found in the behavioral experiment.

**Figure 6:**
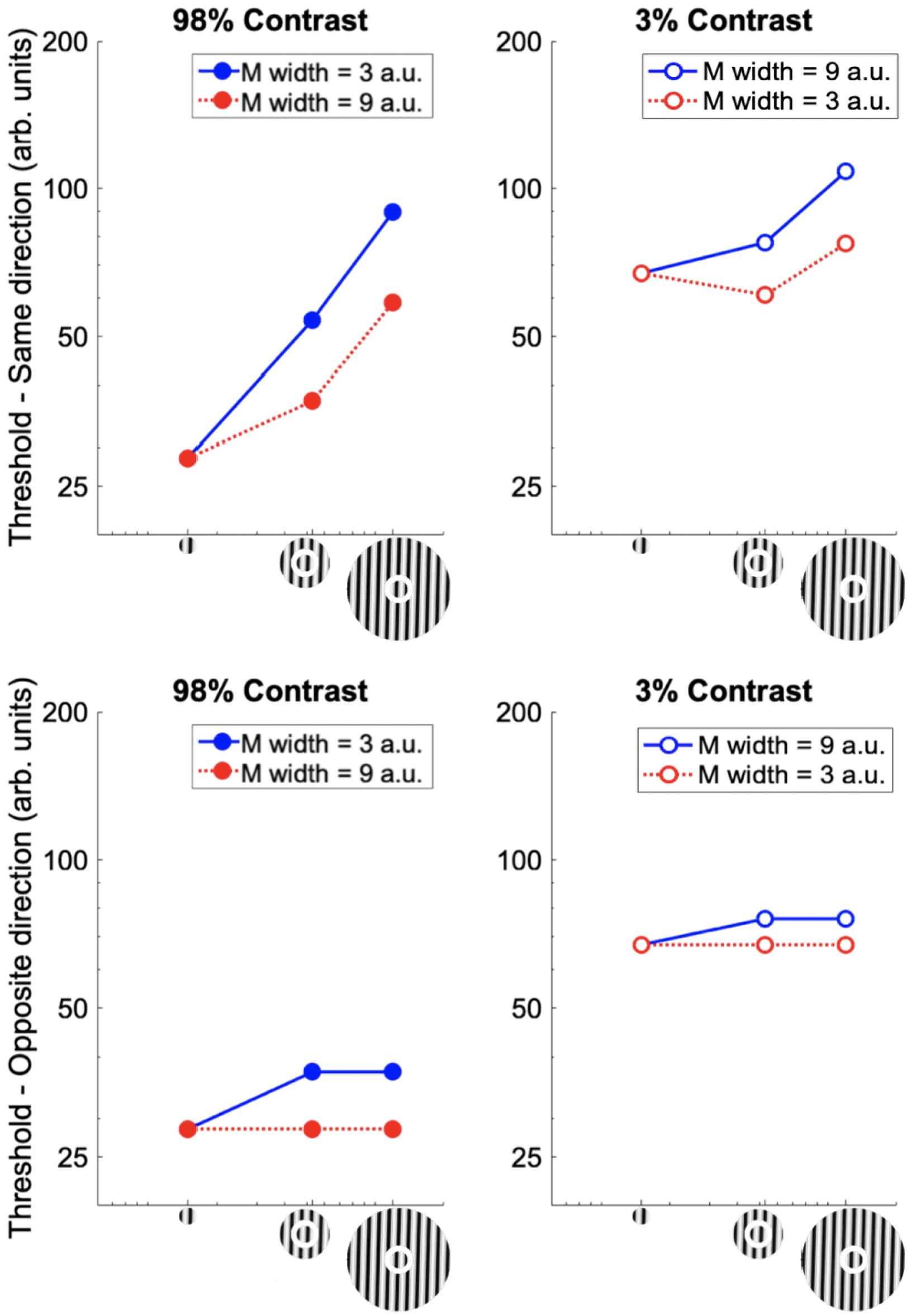
Model results. Predictions of normalization model for the same-direction trials (A and B), and opposite-direction trials (C and D).

Besides, our model predicts that thresholds increase as the attention field increases not only for the same-direction trials but also for the opposite-direction trials. Furthermore, the model predicts that the increase in surround suppression with a larger attention field is weaker in opposite-direction trials compared to same-direction trials, which is also consistent with the behavioral data. These results argue that the behavioral effect of attention on surround suppression could be arising from the neuronal activity in (h)MT.

## 4 Discussion

Here, we studied how the spatial extent of attention modulates surround suppression in motion perception. In a behavioral experiment, we measured the duration thresholds for discriminating motion direction of foveally presented drifting gratings under wide and narrow spatial attention conditions. Then, by using a computational model that incorporates the spatial extent of attention and response normalization (Reynolds & Heeger, 2009), we sought a link between the behavioral results and possible neural activity patterns. First, we found that increasing the spatial extent of attention evoked stronger surround suppression. Second, the magnitude of the effect increased as the surround size increased for low-contrast stimuli, but did not change for high-contrast stimuli. Third, attention had a weaker effect when the center and surround gratings drifted in opposite directions. Finally, using the divisive normalization model of attention, we showed that the neuronal activity in area hMT may predict the behavioral findings.

### Wider spatial attention leads to stronger surround suppression

We found that surround suppression was significantly stronger under the wide-attention condition compared to the narrow-attention condition. This is consistent with previous studies in literature where the modulatory effect of attention on surround suppression has been commonly demonstrated using a paradigm in which observers are asked to attend versus ignore a static stimulus (Flevaris & Murray, 2015a; Herrmann et al., 2010; Itthipuripat, Garcia, Rungratsameetaweemana, Sprague, & Serences, 2014; Sundberg, Mitchell, & Reynolds, 2012; Williford & Maunsell, 2006). However, unlike in previous studies, here we investigated the effect of the spatial extent of attention rather than mere presence or absence of spatial attention. Hence,to the best of our knowledge, our results are the first to systematically demonstrate that surround suppression in motion perception becomes stronger with a wider attention field.

Note that, even though there was a difference in the strength of surround suppression between the wide and narrow attention conditions for small-surround low-contrast stimuli, suppression indexes (*SI*) for those conditions did not reach significance (*SI* marginally significant for wide attention condition, *p* = 0.008; *α_corr_* = 0.00625). This is not surprising, and it is consistent with previous studies reporting surround facilitation with low-contrast and relatively small stimuli (Er et al., 2020; Schallmo et al., 2018; Tadin et al., 2003). Moreover, the pattern of our results, that is the existence of surround suppression in high but not low contrast small stimuli, is also consistent with the response characteristics of neurons when a stimulus is presented in their non-classical receptive fields (RF) (Angelucci & Bressloff, 2006; Shushruth et al., 2013).

### Spatial attention interacts with size and contrast of the stimulus

We found that *SI*, on the average, increases with size more strongly under the wide compared to narrow attention condition. Assessing this effect further, we found that the difference in *SI*s between the narrow and wide attention conditions increases with size for low contrast stimuli, but does not change for high contrast stimuli. This pattern of results may be partly related to the non-classical RF characteristics discussed in the previous paragraph.

It has been known that spatial attention affects contrast-response function and surround suppression at the perceptual level, but how it interacts with other contextual factors such as size and contrast of the stimulus was not systematically studied before in human motion perception. Our results here provide further knowledge to close this gap in the literature.

### Spatial attention has a weaker effect when center and surround move in opposite direction

We found that the attention effect (the difference in *SI*s under the wide and narrow attention conditions) was smaller when the center and surround drifted in opposite directions compared to when they drifted in the same direction. This shows that the effect of attention on surround suppression under the same-direction condition is not an artifact of the task demand. We reason as follows. At the single-unit level surround suppression is reduced or eliminated when surround and center move in opposite directions (Allman et al., 1985; Born & Tootell, 1992; Cavanaugh et al., 2002b; Kastner et al., 1995; Lamme, 1995). Thus, if the attention effect we find in the same direction trials is truly related to the surround suppression mechanisms, then it should be larger than that under the opposite direction condition. If the effect ensues as a result of the higher demand of the second task in the wide attention condition, however, then we would observe no difference between the same- and opposite-direction conditions. Our results show that the effect is stronger under the same-direction condition compared to different direction condition, and addresses this possible task-demand artifact confound.

There is limited information in literature on the behavioral effects of surround suppression when center and surround move in opposite directions (Moutsiana, Field, & Harris, 2011; Paffen, Alais, & Verstraten, 2005a; Paffen, van der Smagt, te Pas, & F.Verstraten, 2005b; Tadin, Paffen, Blake, & Lappin, 2008). For example, Paffen et al. (2005b) found that large, high-contrast surround facilitated opposite-direction motion perception in the center, but this effect was not present for small- or low-contrast surrounds. This is partly consistent with our findings. Namely we also found that for opposite-direction trials, suppression is more prominent in low-contrast trials and it is reduced for high-contrast trials, especially for large-surround conditions. Neurophysiological studies, on the other hand, have shown that attention affects the contrast-response functions when two different motion stimuli, one moving in the preferred one in the opposite direction, are presented simultaneously within the RF of a neuron (Lee & Maunsell, 2010a, 2010b; Martinez-Trujillo & Treue, 2002). To the best of our knowledge, however, our study is the first to demonstrate how attention affects surround suppression when center and surround drift in opposite directions.

### Activity of hMT neurons may explain the behavioral effect

Using the normalization model of attention (Reynolds & Heeger, 2009) and parameters derived from literature, we showed that the neural activity in area hMT may lead to the perceptual effect. This supports the hypothesis that the behavioral manifestation of surround suppression in motion perception originates in hMT (Tadin et al., 2003). Our simulation results are consistent with several studies in literature where hMT activity, measured with fMRI, was shown to be consistent with the behavioral effect (Er et al., 2020; Schallmo et al., 2018; Turkozer et al., 2016). Furthermore, our results show that a simple computational model can qualitatively predict the behavioral effects of surround suppression for a wide variety of factors, including contrast, size, attention, and direction of motion, thus paves the way for future studies.

### Clinical populations, surround suppression and spatial attention

Surround suppression is weaker in certain clinical conditions including schizophrenia (Tadin et al., 2006), major depressive disorder (Golomb et al., 2009) and autism spectrum disorder (ASD) (Foss-Feig et al., 2013; Schallmo et al., 2020; Sysoeva, Galuta, Davletshina, Orekhova, & Stroganova, 2017). The reasons underlying this interesting phenomenon are still hotly debated. Previously a weaker GABAergic system in the patients was put forward to explain the findings (Tadin et al., 2006), but this hypothesis was not supported with more recent MR spectroscopy studies (Schallmo et al., 2018). On the other hand, it is well known that attention mechanisms are affected under many clinical disorders (Clayton, Richards, & Edwards, 1999; Habermann-Paelecke, Pohl, & Leplow, 2005; Kreither et al., 2017; Luck & Gold, 2008). It is possible that abnormal attention leads to abnormal surround suppression in these patients. In line with this conjecture, in a recent study Schallmo et al. (2020) found weaker suppression in ASD patients and suggested that this could be because of narrower spatial attention. Thus, we contend that attention may provide an alternative explanation for the abnormal surround suppression found in clinical populations.

## 5 Conclusion

To conclude, we found that wider spatial attention leads to stronger surround suppression in motion perception. Through computer simulations, we showed that the activity patterns of hMT neurons might cause this effect. Finally, we argue that abnormal attention mechanisms may lead to the abnormal surround suppression observed in some clinical populations.

## Notes

### Competing Interest Statement

The authors have declared no competing interest.

## References

Allman, J., Miezin, F., & McGuinness, E. (1985). Direction- and velocity-specific responses from beyond the classical receptive field in the middle temporal visual area (mt). Perception, 14, 105–126. doi: https://doi.org/10.1068/p140105

Angelucci, A., Bijanzadeh, M., Nurminen, L., Federer, F., Merlin, S., & Bressloff, P. C. (2017). Circuits and mechanisms for surround modulation in visual cortex. Annual Review of Neuroscience, 40, 425–451. doi: 10.1146/annurev-neuro-072116-031418

Angelucci, A., & Bressloff, P. C. (2006). The contribution of feedforward, lateral and feedback connections to the classical receptive field center and extra-classical receptive field surround of primate v1 neurons. Progress in Brain Research, 154, 93–120. doi: https://doi.org/10.1016/S0079-6123(06)54005-1

Anton-Erxleben, K., Stephan, V. M., & Treue, S. (2009). Attention reshapes center-surround receptive field structure in macaque cortical area mt. Cerebral Cortex, 19, 2466–478. doi: 10.1093/cercor/bhp002

Borghuis, B., Tadin, D., Lankheet, M., Lappin, J., & van de Grind, W. (2019). Temporal limits of visual motion processing: Psychophysics and neurophysiology. Vision, 3 (1), 5. doi: 10.3390/vision3010005..

Born, R. T., & Tootell, R. B. H. (1992). Segregation of global and local motion processing in primate middle temporal visual area. Nature, 357, 497–499. doi: 10.1038/357497a0

Carandini, M., & Heeger, D. J. (2012). Normalization as a canonical neural computation. Nature Reviews Neuroscience, 13 (1), 51–62. doi: https://doi.org/10.1038/nrn3136

Cavanaugh, J. R., Bair, W., & Movshon, J. A. (2002a). Nature and interaction of signals from the receptive field center and surround in macaque v1 neurons. Journal of Neurophysiology, 88 (4), 2530–2546. doi: https://doi.org/10.1152/jn.00692.2001

Cavanaugh, J. R., Bair, W., & Movshon, J. A. (2002b). Selectivity and spatial distribution of signals from the receptive field surround in macaque v1 neurons. Journal of Neurophysiology, 88, 2547–2556. doi: 10.1152/jn.00693.2001

Clayton, I. C., Richards, J. C., & Edwards, C. J. (1999). Selective attention in obsessive-compulsive disorder. Journal of Abnormal Psychology, 108 (1), 171–175. doi: https://doi.org/10.1037//0021-843x.108.1.171

Er, G., Pamir, Z., & Boyaci, H. (2020). Distinct patterns of surround modulation in v1 and hmt+. NeuroImage, 220, 117084. doi: https://doi.org/10.1016/j.neuroimage.2020.117084

Flevaris, A. V., & Murray, S. O. (2015a). Attention determines contextual enhancement versus suppression in human primary visual cortex. The Journal of Neuroscience, 35 (35), 12273–12280. doi: 10.1523/JNEUROSCI.1409-15.2015

Flevaris, A. V., & Murray, S. O. (2015b). Feature-based attention modulates surround suppression. Journal of Vision, 15 (1), 1–11. doi: 10.1167/15.1.29

Foss-Feig, J. H., Tadin, D., Schauder, K. B., & Cascio, C. J. (2013). A substantial and unexpected enhancement of motion perception in autism. The Journal of Neuroscience, 33 (19), 8243–8249. doi: https://doi.org/10.1523/JNEUROSCI.1608-12.2013

Freeman, E., Sagi, D., & Driver, J. (2001). Lateral interactions between targets and flankers in low-level vision depend on attention to the flankers. Nature Neuroscience, 4, 1032–1036. doi: 10.1038/nn728

Golomb, J. D., McDavitt, J. R. B., Ruf, B. M., Chen, J. I., Saricicek, A., Maloney, K. H., … Bhagwagar, Z. (2009). Enhanced visual motion perception in major depressive disorder. Journal of Neuroscience, 29 (28), 9072–9077. doi: https://doi.org/10.1523/JNEUROSCI.1003-09.2009

Habermann-Paelecke, Y., Pohl, J., & Leplow, B. (2005). Attention and executive functions in remitted major depression patients. Journal of Affective Disorders, 89 (1-3), 125–135. doi: https://doi.org/10.1016/j.jad.2005.09.006

Heeger, D. J. (1992). Normalization of cell responses in cat striate cortex. Visual Neuroscience, 9 (2), 181–197. doi: 10.1017/S0952523800009640

Herrmann, K., Montaser-Kouhsari, L., Carrasco, M., & Heeger, D. J. (2010). When size matters: attention affects performance by contrast or response gain. Nature Neuroscience, 13 (12), 1554–1559. doi: https://doi.org/10.1038/nn.2669

Ito, M., & Gilbert, C. D. (1999). Attention modulates contextual influences in the primary visual cortex of alert monkeys. Neuron, 22, 593–604. doi: https://doi.org/10.1016/S0896-6273(00)80713-8

Itthipuripat, S., Garcia, J. O., Rungratsameetaweemana, N., Sprague, T. C., & Serences, J. T. (2014). Changing the spatial scope of attention alters patterns of neural gain in human cortex. Journal of Neuroscience, 34 (1), 112–123. doi: https://doi.org/10.1523/JNEUROSCI.3943-13.2014

Kastner, S., Nothdurft, H. C., & Pigarev, I. N. (1995). Neuronal correlates of pop-out in cat striate cortex. Vision Research, 37, 371–376. doi: https://doi.org/10.1016/S0042-6989(96)00184-8

Kingdom, F. A. A., & Prins, N. (2010). Psychophysics: A practical introduction. London: Academic Press: an imprint of Elsevier.

Kreither, J., Lopez-Calderon, J., Leonard, C. J., Robinson, B. M., Ruffle, A., Hahn, B., … Luck, S. J. (2017). Electrophysiological evidence for hyperfocusing of spatial attention in schizophrenia. Journal of Neuroscience, 37 (14), 3813–3823. doi: https://doi.org/0.1523/JNEUROSCI.3221-16.2017

Lamme, V. A. F. (1995). The neurophysiology of figure-ground segregation in primary visual cortex. Journal of Neuroscience, 15, 1605–1616. doi: https://doi.org/10.1523/JNEUROSCI.15-02-01605.1995

Lee, J., & Maunsell, J. H. R. (2010a). Attentional modulation of mt neurons with single or multiple stimuli in their receptive fields. Journal of Neuroscience, 30 (8), 3058–3066. doi: https://doi.org/10.1523/JNEUROSCI.3766-09.2010

Lee, J., & Maunsell, J. H. R. (2010b). The effect of attention on neuronal responses to high and low contrast stimuli. Journal of Neurophysiology, 104, 960–971. doi: doi:10.1152/jn.01019.2009

Luck, S. J., & Gold, J. M. (2008). The construct of attention in schizophrenia. Biological Psychiatry, 64 (1), 34–39. doi: https://doi.org/10.1016/j.biopsych.2008.02.014

Martinez-Trujillo, J. C., & Treue, S. (2002). Attentional modulation strength in cortical area mt depends on stimulus contrast. Neuron, 35, 2. doi: https://doi.org/10.1016/S0896-6273(02)00778-X

Maunsell, J. H. R. (2015). Neuronal mechanisms of visual attention. Annual Review of Vision Science, 1, 373–391. doi: 10.1146/annurev-vision-082114-035431

Moutsiana, C., Field, D. T., & Harris, J. P. (2011). The neural basis of centre-surround interactions in visual motion processing. PLoS ONE, 6 (7), e22902. doi: 10.1371/journal.pone.0022902

Paffen, C. L. E., Alais, D., & Verstraten, F. A. (2005a). Center–surround inhibition deepens binocular rivalry suppression. Vision Research, 45, 2642–2649. doi: 10.1016/j.visres.2005.04.018

Paffen, C. L. E., van der Smagt, J., Maarten, te Pas, S., & F. Verstraten. (2005b). Center-surround inhibition and facilitation as a function of size and contrast at multiple levels of visual motion processing. Journal of Vision, 5, 571–578. doi: 10.1167/5.6.8

Petrov, Y., Carandini, M., & McKee, S. (2005). Two distinct mechanisms of suppression in human vision. Journal of Neuroscience, 25 (38), 8704–8707. doi: https://doi.org/10.1523/JNEUROSCI.2871-05.2005

Reynolds, J. H., & Chelazzi, L. (2004). Attentional modulation of visual processing. Annual Review of Neuroscience, 27 (1), 611–647. doi: 10.1146/annurev.neuro.26.041002.131039

Reynolds, J. H., & Heeger, D. J. (2009). The normalization model of attention. Neuron, 61 (2), 168–185. doi: https://doi.org/10.1016/j.neuron.2009.01.002

Roberts, M., Delicato, L. S., Herrero, J., Gieselmann, M. A., & Thiele, A. (2007). Attention alters spatial integration in macaque v1 in an eccentricity-dependent manner. Nature Neuroscience, 10, 1483–1491. doi: https://doi.org/10.1016/S0896-6273(00)80713-8

Schallmo, M.-P., Kale, A. M., Flevaris, A. V., Brkanac, Z., Edden, R. A., Bernier, R. A., & Murray, S. O. (2018). Suppression and facilitation of human neural responses. eLife, 7, 1–23. doi: https://doi.org/10.7554/eLife.30334.001

Schallmo, M.-P., Kolodny, T., Kale, A. M., Millin, R., Flevaris, A. V., Edden, R. A., … Murray, S. O. (2020). Weaker neural suppression in autism. Nature Communications, 11, 2675. doi: https://doi.org/10.1038/s41467-020-16495-z

Shushruth, S., Mangapathy, P., Ichida, J. M., Bressloff, P. C., Schwabe, L., & Angelucci, A. (2012). Strong recurrent networks compute the orientation tuning of surround modulation in the primate primary visual cortex. The Journal of Neuroscience, 32, 308–321. doi: https://doi.org/10.1523/JNEUROSCI.3789-11.2012

Shushruth, S., Nurminen, L., Bijanzadeh, M., Ichida, J. M., Vanni, S., & Angelucci, A. (2013). Different orientation tuning of near- and far-surround suppression in macaque primary visual cortex mirrors their tuning in human perception. The Journal of Neuroscience, 33 (1), 106–119. doi: 10.1523/JNEUROSCI.2518-12.2013

Sundberg, K. A., Mitchell, J. F., & Reynolds, J. H. (2009). Spatial attention modulates center-surround interactions in macaque visual area v4. Neuron, 61 (6), 952–963. doi: 10.1016/j.neuron.2009.02.023

Sundberg, K. A., Mitchell, J. F., & Reynolds, J. H. (2012). Attention influences single unit and local field potential response latencies in visual cortical area v4. The Journal of Neuroscience, 32 (45), 16040–16050. doi: 10.1523/JNEUROSCI.0489-12.2012

Sysoeva, O. V., Galuta, I. A., Davletshina, M. S., Orekhova, E. V., & Stroganova, T. A. (2017). Abnormal size-dependent modulation of motion perception in children with autism spectrum disorder (asd). Frontiers in Neuroscience, 11, 164. doi: https://doi.org/10.3389/fnins.2017.00164

Tadin, D. (2015). Suppressive mechanisms in visual motion processing: From perception to intelligence. Vision Research, 115, 58–70. doi: https://doi.org/10.1016/j.visres.2015.08.005

Tadin, D., Kim, J., L. Doop, M., Gibson, C., Lappin, J. S., Blake, R., & Park, S. (2006). Weakened center-surround interactions in visual motion processing in schizophrenia. Journal of Neuroscience, 26 (44), 11403–11412. doi: https://doi.org/10.1523/JNEUROSCI.2592-06.2006

Tadin, D., Lappin, J. S., Gilroy, L. A., & Blake, R. (2003). Perceptual consequences of centre-surround antagonism in visual motion processing. Nature, 424, 312–315. doi: https://doi.org/10.1038/nature01800

Tadin, D., Paffen, C. L. E., Blake, R., & Lappin, J. S. (2008). Contextual modulations of center-surround interactions in motion revealed with the motion aftereffect. Journal of Vision, 8 (7), 1–11. doi: https://doi.org/10.1167/8.7.9

Turkozer, H. B., Pamir, Z., & Boyaci, H. (2016). Contrast affects fmri activity in middle temporal cortex related to center–surround interaction in motion perception. Frontiers in Psychology, 7, 1–8. doi: 10.3389/fpsyg.2016.00454

Williford, T., & Maunsell, J. H. (2006). Effects of spatial attention on contrast response functions in macaque area v4. Journal of Neurophysiology, 96 (1), 40–54. doi: 10.1152/jn.01207.2005

Womelsdorf, T., Anton-Erxleben, K., Pieper, F., & Treue, S. (2006). Dynamic shifts of visual receptive fields in cortical area mt by spatial attention. Nature Neuro-science, 9, 1156–1160. doi: 10.1167/15.1.29

Xing, J., & Heeger, D. J. (2001). Measurement and modeling of center-surround suppression and enhancement. Vision Research, 41, 571–583. doi: https://doi.org/10.1016/S0042-6989(00)00270-4

